# An IL-1, IL-17, and IL-22 cytokine circuit controls vulvovaginal candidiasis independently of estrogen

**DOI:** 10.1101/2025.08.25.671995

**Authors:** Bianca M. Coleman, Melissa E Cook, Md. Robin Khan, Amanda K. Vogel, Anthony J. Wells, Jian Miao, Shachi P. Vyas, Tiffany C. Taylor, Felix E.Y. Aggor, Nicole O. Ponde, Ipsita Dey, Henry Zou, Eldin Jašarević, Brian M. Peters, Sarah L. Gaffen

## Abstract

Vulvovaginal candidiasis (VVC) affects >75% of women, with considerable morbidity and high medical cost burden. While Type 17 cytokines (IL-17, IL-22) are critical for oral and dermal immunity to *C. albicans*, their role in VVC has been less clear. Th17 gene signatures are potently upregulated in VVC, yet impairment of individual Th17 components (IL-17A, IL-17R subunits, IL-22) does not worsen disease. Rather, estrogen activity is tightly linked to VVC, leading to a paradigm that hormonal pathways rather than immune defense, dominate susceptibility. Here, we reveal a previously unappreciated role for IL-1/Type 17 in VVC that operates independently of estrogenic hormones. In contrast to mice lacking IL-17A, IL-17RA, IL-22, or IL-22R individually, mice lacking IL-17RA and IL-22RA1 together (*Il17raIl22ra1*^-/-^) exhibited high fungal loads and exacerbated tissue damage and inflammation. In human vulvar epithelial cells, IL-17 and IL-22 drive synergistic signaling. IL-1R signaling but surprisingly not IL-23 was upstream of this response. *Il17raIl22ra1*^-/-^ mice expressed high IL-1β yet did not control disease, indicating that IL-1 is upstream but not downstream of Type 17 responses. Unexpectedly, Type 17-dependent control occurred in the absence of exogenous estrogen administration and persisted even when estrus was prevented by progesterone treatment. Collectively, these data indicate that susceptibility to VVC is driven not only by estrogen sensitization but through combinatorial loss of IL-17 and IL-22.

**Graphical Abstract:** 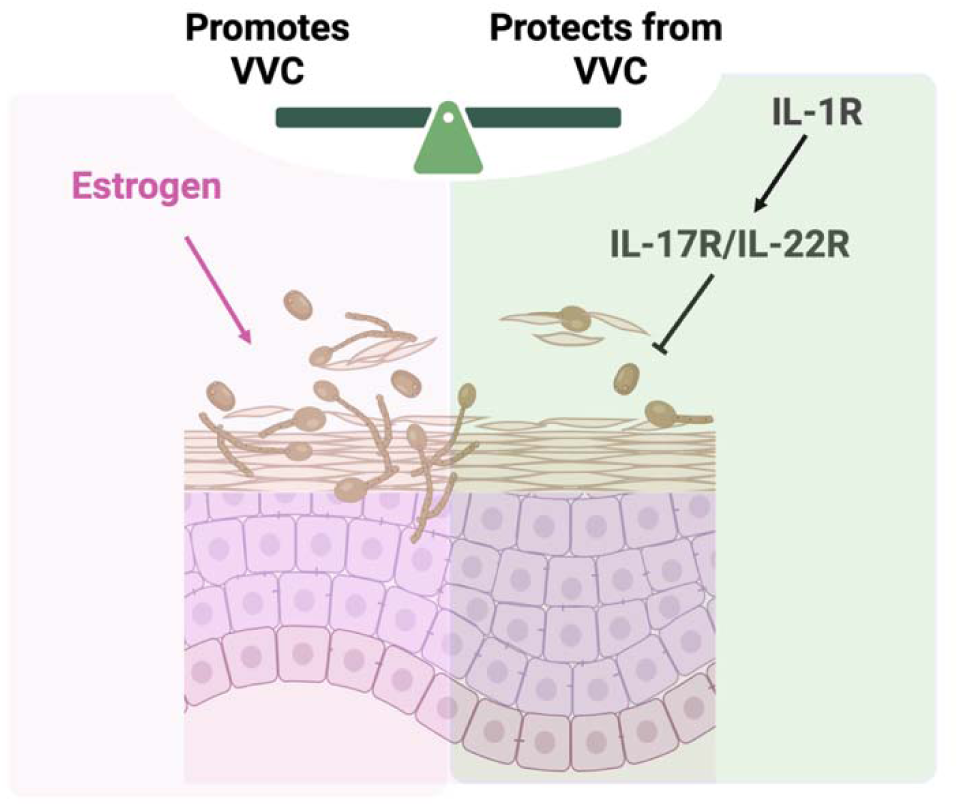

## Introduction

The commensal pathobiont *Candida albicans* colonizes healthy individuals and is a designated World Health Organization critical priority pathogen [1]. Vulvovaginal candidiasis (VVC) and oropharyngeal candidiasis (OPC) are the most common manifestations of candidiasis, while systemic infections are rarer though deadly [2]. The tissue-specific immune mechanisms of host defense to *C. albicans* are surprisingly distinct. OPC arises primarily from T cell immunocompromise (e.g., HIV/AIDS), IL-17/Th17 deficiency, or impairments in oral immunity (e.g., head-neck irradiation or xerostomia) [2, 3]. In contrast, VVC occurs in at least 75% of otherwise healthy women. For many, VVC occurs more than 3 times per year, imposing a major detriment on quality of life and mental health [4–7]. Contributing factors to VVC susceptibility are microbiota abundance and composition, vaginal pH, behavioral factors, diabetes, and elevation in estrogenic hormones that establish a permissive environment for infection [7]. Although often treatable with over-the-counter antifungal medications, drug resistance is an increasing problem and medical costs are significant [4, 7].

Typically, VVC is considered a condition of immunopathology, as symptomatic infection correlates with high polymorphonuclear leukocytes (PMN) in the vaginal lumen accompanied by elevated expression of inflammatory cytokines and markers of epithelial damage [4, 8–11]. Although elevated, neutrophils in the vaginal microenvironment do not adequately facilitate fungal clearance [12]. Unique among manifestations of candidiasis, VVC is closely linked to estradiol, the most potent form of estrogen. Disease is rare in pre-pubescent and post-menopausal individuals (who have low estradiol) and high in patients receiving estrogen-based contraceptives, hormone replacement therapy, or during high estrogenic phases of pregnancy. In rodents, exogenous estradiol administration is required to establish persistent vaginal *Candida* colonization [13–15]. Estrogen is reported to act on both host and fungal cells to establish a permissive environment for VVC, though the precise mechanisms are still not well elucidated [4, 7].

Nonetheless, adaptive immune responses contribute to control of VVC, though the role of the Type 17 pathway is not as clear as in oral and dermal candidiasis. Infections with avirulent *C. albicans* strains protect mice from subsequent infections, as does immunization with recombinant *C. albicans* cell wall antigens [4, 16]. In humans, a Phase Ib/IIa clinical trial of a *Candida* vaccine, patients generated *Candida-*specific Th1 and Th17 adaptive responses, increased IgG specific antibodies, and a subset of patients with recurrent VVC (RVVC) showed significantly delayed time to recurrence [17–20]. *C. albicans* drives IL-17 production in human PBMCs [21], and induces a pan-fungal Th17 cell response in humans [22]. Th17 gene signatures are elicited in human and murine vaginal tissue during *C. albicans* infection [23–25], and VVC is a statistically significant, albeit modest, adverse event associated with anti-IL-17 biologic therapy [26]. In mice, long term control requires T cells [27], and vaccination against *C. albicans* generates Th17 responses [16]. Halofuginone, an aminoacyl-tRNA synthetase inhibitor that restrains Th17 cells [28, 29], worsens vaginal candidiasis in mice [23]. However, in contrast to OPC, infection with HIV/AIDS or loss of function mutations in IL-17RA or other Type 17 cytokine genes are not linked to VVC incidence in humans [4, 30]. Similarly in mice, deletion of individual genes within the Th17 pathway (*Il17a, Il17ra*, *Il22, Act1)* does not increase susceptibility to VVC [6, 24, 31]. Thus, the accepted paradigm is that VVC is distinct among manifestations of candidiasis in not being subject to control by Th17 cytokines.

Here, we provide evidence that Type 17 pathway does provide immune control of VVC, potentially reconciling these conflicting observations in the field. While loss of IL-17RA/IL-17A or IL-22/IL-22R do not render mice susceptible to VVC ([24, 31], this report) we show that mice with a combined loss of both cytokine pathways (*Il17ra^-/-^Il22ra1^-/-^)* exhibit not only high fungal colonization but also elevated tissue damage and inflammation. In a human vaginal epithelial cell line, IL-17 and IL-22 synergistically upregulated many canonical antifungal immune genes. IL-1R- and IL-1β-deficient mice displayed similar increased susceptibility to VVC. However, high levels of IL-1β were found in the absence of IL-17/22 signaling yet were insufficient to control disease, thus indicating that IL-1 receptor signaling is upstream but not downstream of the Th17 axis. Moreover, Type 17 control of VVC was independent of estrogen-driven susceptibility, as high fungal loads occurred in IL-17R/IL-22R-deficient and IL-1R-deficient mice without exogenous estrogen treatment. Susceptibility persisted even when estrus was suppressed with progesterone, a hormone known to promote fungal clearance. Finally, when estrogen treatment was combined with IL-17R/IL-22R-deficiency, there was far higher disease burden than in either condition alone. Together, these results reveal an underlying estrogen-independent contributor to immunity to *C. albicans* in the vaginal mucosa driven by Type 17 cytokines and IL-1R signaling.

## Results

### IL-1 receptor signaling mediates immunity to VVC and activates Type 17 cytokines

Several studies have implicated the inflammasome and IL-1 cytokines in VVC [24, 32–36], but how this pathway functions in disease is not well defined. To determine how IL-1R signals function in VVC, *Il1r*^-/-^ mice lacking the receptor for IL-1α and IL-1β were subjected to a standard model of VVC [15] (**Fig 1a)**. Mice were administered 17-β-estradiol (E2) on days -3 and 4 relative to vaginal administration with *C. albicans*. Fungal loads in vaginal lavage fluid (VLF) were assessed by CFU enumeration on days 3 and 7 post-infection. *Il1r*^-/-^ mice showed approximately 8-fold higher vaginal colonization of *C. albicans* compared to WT at both early and late time points (**Fig 1b**), confirming a protective role for IL-1R signaling in VVC. Similarly, mice lacking IL-1β (*Il1b*^-/-^) showed higher CFUs than WT (**Fig. 1c**). Histologically, *Il1r*^-/-^ vaginal tissue exhibited more *C. albicans* hyphae and evidence of mucosal damage compared to controls (**Fig 1d**). Consistent with this, levels of lactate dehydrogenase levels (LDH, released during hyphal-mediated epithelial destruction by candidalysin [5, 37, 38]) in VLF were elevated in *Il1r*^-/-^ and *Il1b*^-/-^ mice upon infection (**Fig 1e**). There was not a significant increase in LDH levels at baseline in *Il1r^-/-^* mice (**Fig S1A**). PMN infiltration to the vaginal lumen, considered a marker of immune pathology, trended lower on day 3 in *Il1r*^-/-^ mice and was significantly reduced on day 7 (**Fig 1f).** Unexpectedly, in IL-1β-deficient mice, PMNs were elevated at both time points (**Fig 1f**), indicating a disconnect between IL-1R-mediated control of fungal loads versus PMNs.

**Fig. 1.**
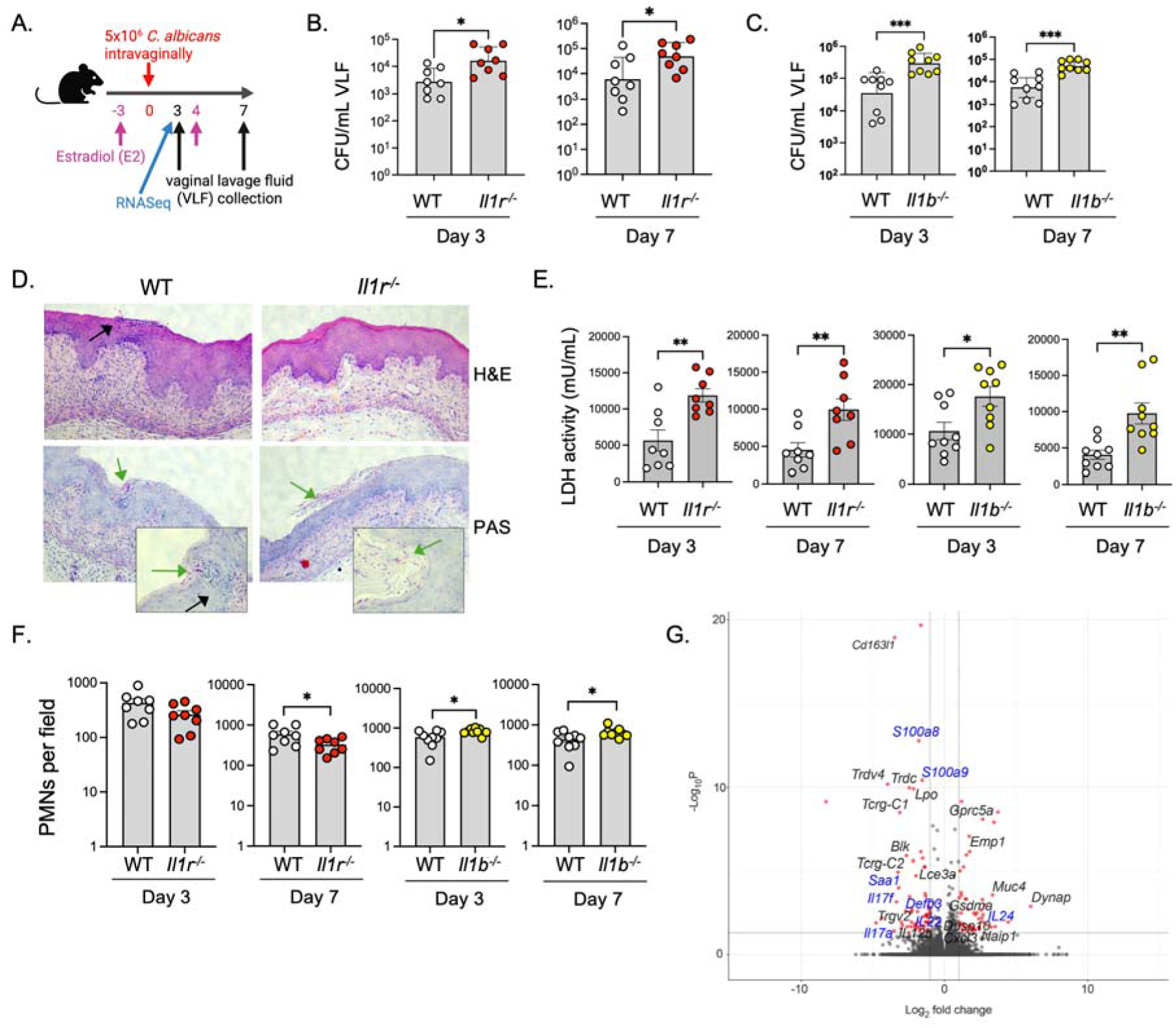
**IL-1R signals control vaginal immunity to VVC through IL-1**β **a.** VVC model **b, c.** Mice were administered E2 on day -3, and on day 4 were infected vaginally with C. albicans. Fungal burdens in WT (C57BL/6, n=8-9), Il1r^-/-^ (n=8) or Il1b^-/-^ (n=9) mice were assessed by CFU enumeration on day 3 and 7. Geometric mean ± geometric SD and Mann-Whitney U test. **d.** Representative vaginal histology (day 3) stained with H&E or Periodic acid- Schiff (PAS). Green arrows denote C. albicans hyphae. Black arrows indicate PMNs **e.** LDH levels in VLF (mU/mL). Mean+SEM, analyzed by t-test with Welch’s correction. **f.** PMN counts per field in VLF. Mean ± SEM, analyzed by two-sided Student’s t-test. **g.** Differentially expressed genes in vaginal tissue of WT compared to Il1r^-/-^ mice at day 3 (adjusted p < 0.05, Student’s t-test and Benjamini and Hochberg’s analysis). Genes associated with the Type 17 response indicated in blue.

The mechanisms by which the IL-1R pathway drives antifungal host defense are not well defined. We subjected vaginal mucosal tissue from infected *Il1r^-/-^* mice to bulk RNASeq analysis on day 3 post-infection (**Fig 1g**). *Il1r^-/-^* mice showed impaired expression of Type 17 cytokines such as *Il17a, Il17f,* and *Il22*. Also reduced in *Il1r^-/-^*mice were signature downstream gene targets associated with the Type 17 response and IL-17 signaling, for example *Defb3, Saa1, S100a8/a9* [39, 40] (**Fig 1g**). These findings are in line with findings that IL-1 directs differentiation and activation of Type 17 cells [41] and that expression of Type 17 genes correlates with VVC [23–25].

### Cooperative signals from IL-17 and IL-22 but not IL-23 control C. albicans in the vaginal tract

These results prompted us to reexamine the possibility that the Type 17 axis participates in VVC immunity. In addition to IL-1, IL-23 is a key upstream activator of Type 17 cell cytokine production and is required for immunity to oral, dermal and systemic candidiasis [39, 42–45]. Given the impaired expression of Type 17 cytokine transcripts in *Il1r*^-/-^ vaginal tissue, including *Il12b* (IL-12p40, the shared subunit of IL-12 and IL-23), we asked whether IL-23 was also required for *C. albicans* colonization in VVC. In contrast to *Il1r^-/-^*mice and consistent with a prior study [31], mice lacking IL-23 did not exhibit elevated fungal loads in E2-induced VVC (**Fig 2a**). Neither *Il17a, Il17f* mRNA nor IL-17A protein levels were statistically lower in the vaginal mucosa of infected *Il23^-/-^* mice. *Il1b* mRNA and IL-1β were also unchanged (**Fig 2b**). Thus, IL-1 rather than IL-23 appears to be the central regulator of the IL-17 response in vaginal candidiasis.

**Figure 2.**
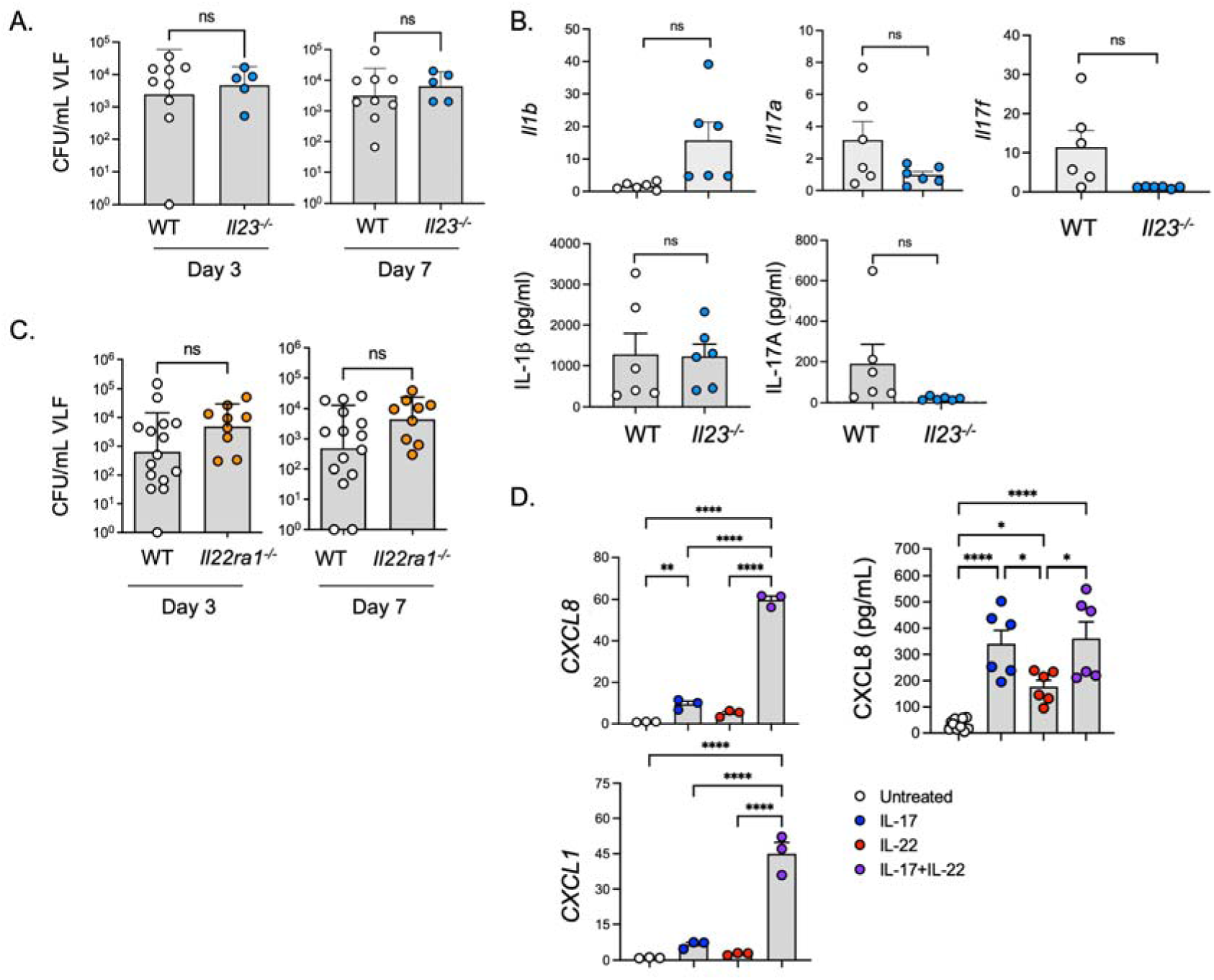
Combined loss of IL-17R and IL-22R signaling increases VVC susceptibility. **a.** Fungal loads in VLF of E2-treated Il23a^-/-^ mice (n=5-9) mice on days 3 and 7 post-infection. geometric mean ± geometric SD, two-tailed unpaired Mann-Whitney test. **b.** Cytokine mRNA and protein levels in vaginal tissue of Il23a^-/-^ mice were assessed on day 3. Mean+SEM, t-test with Welch’s correction. **c.** Fungal loads in VLF from Il22ra^-/-^ (n=9-15) mice. geometric mean ± geometric SD, two-tailed unpaired Mann-Whitney test **d.** A-431 vulvar epithelial cells were treated with human IL-17 or IL-22 for 60 mins. Left: CXCL1 and CXCL8 mRNA assessed by qPCR relative to GAPDH and normalized to untreated control (0 minutes). mean+SEM, analyzed by ANOVA and Kruskal-Wallis test with Dunn’s multiple comparisons. Right: supernatants from A-431 cells treated with IL-17 or IL-22 for 6 h were assessed by ELISA. Mean+SEM, analyzed by one-way ANOVA with Bonferroni’s multiple comparisons test.

Transcriptomic data from infected *Il1r^-/-^* vaginal tissue showed alterations in expression of several cytokines that use the shared IL-22RA1 receptor subunit, including *Il22, Il20* and *Il24* (**Fig 1g**) [46]. We previously showed that loss of IL-22 in mice does not worsen susceptibility to VVC [24], but it was conceivable that other IL-22-family cytokines that were preserved in those mice might serve redundant functions. Therefore, to capture the cumulative impact of the IL-22R-dependent ligands, we evaluated vaginal fungal burdens in *Il22ra1^-/-^* mice. However, mice lacking IL-22RA showed the same susceptibility to VVC as WT counterparts (**Fig 2c**), arguing against additive or redundant impacts of other IL-22R-ligands during vaginal *C. albicans* infection.

Expression of Type 17 signature cytokines (*Il17a, Il17f, Il22*) are among the most significantly induced genes in VVC [24] and were reduced in *Il1r^-/-^* mice (**Fig 1g**). IL-17 synergizes potently with many inflammatory stimuli, and IL-17A/F and IL-22 exhibit cooperative activities in several contexts including OPC [39, 47–52]. Accordingly, while loss of these cytokines individually did not render mice susceptible to VVC, it was possible that their combined activities might be needed for vaginal host defense. A human vulvar epithelial cell line A-431 was treated with human IL-17 and/or IL-22 and known target chemokine, cytokine and antifungal genes were assessed over 6 hours (*CXCL1, CXCL8, IL6, DEFB4A*). Individually, IL-

17 and IL-22 induced these genes modestly, but in combination they showed synergistic induction at the mRNA level (**Fig 2d, Fig S2**). Synergy was less apparent at the protein level, as these cytokines individually induced CXCL8 (IL-8) equivalently (**Fig 2d**).

To examine whether IL-17 and IL-22 exert combinatorial activities *in vivo*, we induced VVC in mice lacking both IL-17RA (which cannot respond to IL-17A, IL-17F, IL-17C and IL-25 [53]) and IL-22RA1 (which are refractory to IL-22, IL-20 and IL-24 [54]). Strikingly, *Il17raIl22ra1^-/-^*mice given E2 and infected with *C. albicans* showed more than 50-fold higher fungal loads compared to WT controls. Higher fungal loads were evident at both days 3 and day 7 (**Fig 3a**). Importantly, this result indicates that E2 sensitization does not confer maximal susceptibility to *C. albicans* colonization, since loss of underlying IL-17/22-dependent signals resulted in much higher fungal loads and tissue damage than with just E2 treatment. Pap smears from *Il17raIl22ra1^-/-^*VLF frequently showed accumulation of *C. albicans* hyphae, which was rarely seen in WT samples (**Fig 3b**). This susceptibility to VVC in *Il17raIl22ra1^-/-^* mice contrasts with results from mice lacking either IL-17RA, IL-22 or IL-22RA1 alone (Ref. [24] and **Fig 2c**), indicating that control of *C. albicans* colonization in the vaginal tract contrasts from the oral mucosa where loss of either cytokine renders mice and humans prone to mucosal candidiasis. Moreover, since Type 17 cytokine-driven signals still protect from VVC even in the presence of E2, these are likely independent routes to VVC.

**Figure 3.**
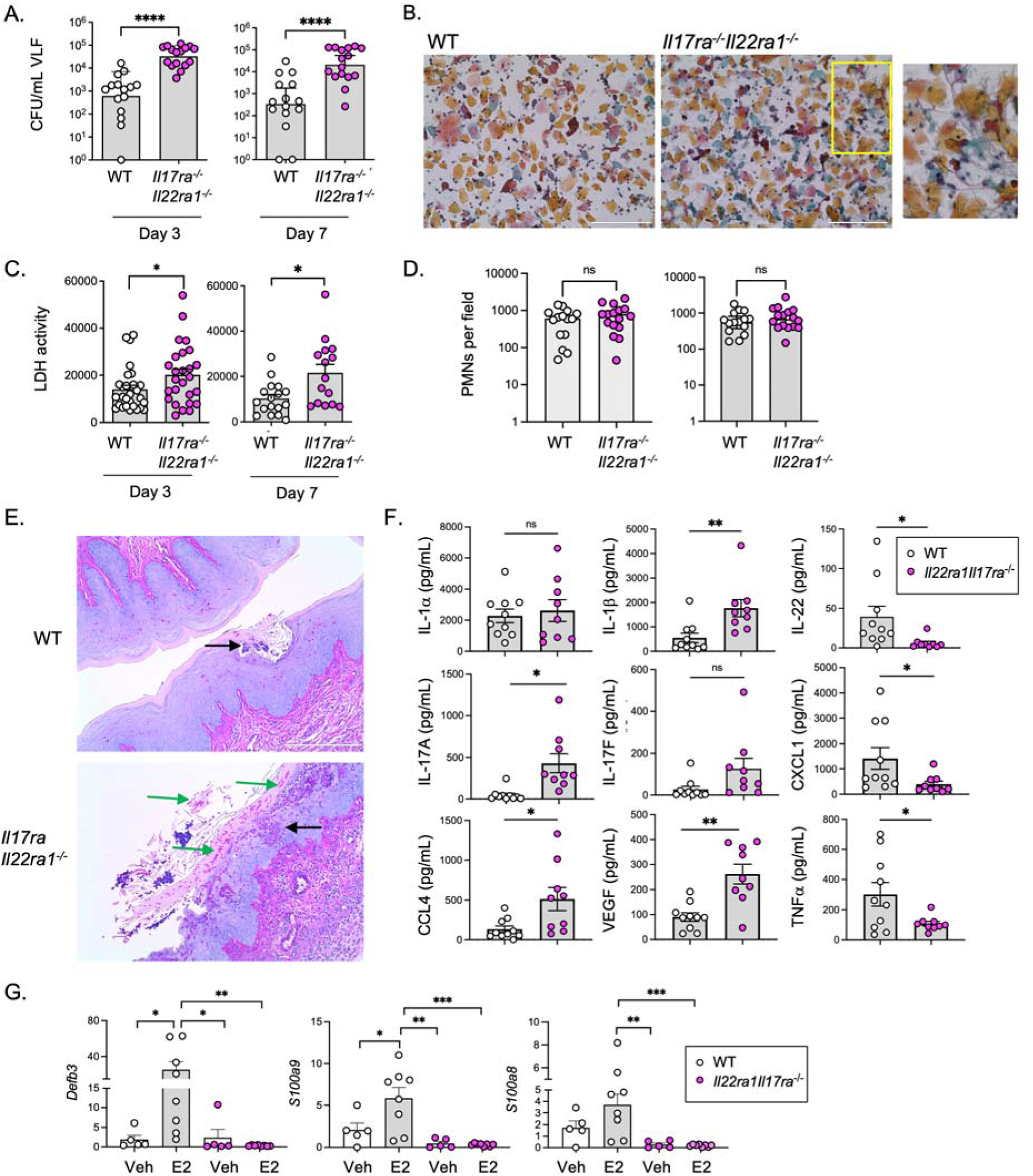
IL-17/22-driven signaling dampens inflammation and tissue damage. **a.** Fungal loads in VLF of E2-treated mice (WT n= 16; Il17raIl22ra1^-/-^ n= 16). mean ± geometric SD, two-tailed unpaired Mann-Whitney test. **b.** Pap smears from WT or Il17raIl22ra1^-/-^ mice. Size bar = 200 µm. Inset: Representative area with visible hyphae. **c.** LDH activity in VLF on days 3 and 7 (WT n= 16; Il17raIl22ra1^-/-^ n= 15). mean ± SEM; unpaired t-test with Welch’s correction **d.** PMNs in VLF from panel A. unpaired t-test with Welch’s correction **e.** Representative PAS and eosin staining of vaginal tract on day 3. Green arrows denote C. albicans hyphae. Black arrows indicate PMNs **f.** Cytokine/chemokine levels determined by Luminex. two-tailed unpaired t-test with Welch’s correction **g.** mRNA levels of indicated genes in vaginal tissue from C. albicans-infected WT (vechicle n=5; E2 n=8) or Il17raIl22ra1^-/-^ (veh n=5; E2 n=8) mice on day 3. Normalized to Gapdh, analyzed by ANOVA and t-test with Tukey’s multiple comparisons.

### IL-17 and IL-22 signaling limits VVC pathology

VVC is characterized by tissue damage and pathology, so fungal colonization alone does constitute a diseased state. We examined LDH levels as a marker of tissue damage, which were elevated nearly 2-fold in *Il17raIl22ra1^-/-^*mice at early and late time points (**Fig 3c)**. LDH correlated with the high fungal burdens in this condition, which interestingly was more pronounced in *Il17raIl22ra1^-/-^* mice than in WT controls (**Fig S3A**). Thus, not only is colonization resistance controlled by IL-17/IL-22 signals, but *C. albicans* is more damaging in this context. However, unlike *Il1r^-/-^* mice, PMN counts in VLF in E2-treated *Il23^-/-^, Il22Ra1^-/-^*and in *Il17raIl22ra1^-/-^* were the same as in WT mice (**Fig 3d, Fig S3B**), despite the high fungal loads that might be expected to provoke a neutrophil response. This observation aligns with findings that PMN function is impaired in the vaginal microenvironment and that neutrophils are insufficient to control fungal loads in VVC [12, 55]. These data also suggest that, since PMNs were impaired in *Il1r*^-/-^ mice but not *Il17raIl22ra1^-/-^* mice (see **Fig 1f**), PMN levels may be governed by an IL-1-dependent but Type 17-independent pathway. Histological evaluation of the *Il17raIl22ra1^-/-^* vagina showed considerable invading hyphae, tissue sloughing and clusters of PMNs along the surface mucosa, indicative of ongoing inflammation and damage (**Fig 3e**).

We also examined VLF levels of 40 cytokines and chemokines associated with inflammation and Type 17 pathways at baseline and during E2-induced VVC (**Fig 3f**). IL-1β and IL-17A were elevated in *Il17raIl22ra1^-/-^*mice; the latter was expected based on studies demonstrating high IL-17A levels in *Il17ra*^-/-^ mice [39, 56], but is also consistent with elevated IL-1β. VEGF and CCL4 were also increased in infected *Il17raIl22ra1^-/-^*VLF, indicative of elevated tissue inflammation and likely driven by elevated fungal loads. In contrast, IL-22, CXCL1, and TNF were lower in *Il17raIl22ra1^-/-^* mice compared to WT (**Fig 3f**). Since diabetes is a risk factor for VVC, we also examined blood glucose levels, which were not different in *Il17raIl22ra1^-/-^* mice compared to WT (**Fig S1C**).

### The IL-1-Type 17 axis restricts fungal burden independently of estrous cycle

VVC correlates with high estradiol phases of life, and all animal models of VVC use estradiol to synchronize cycling and facilitate *C. albicans* colonization [15, 57, 58]. Although the precise mechanisms are not fully elucidated, E2 may play many roles in enabling colonization, including complement evasion, altered cytokine and PRR expression, indirect impairment of neutrophil function and NETosis and stimulation of hyphal growth [13, 59–62]. Most published studies of VVC using gene-targeted mice only examine disease in the presence of E2, so comparatively little is known about estrogen-independent pathways that may contribute to vaginal infection or the impact of cytokine pathways on hormonal fluctuations in the setting of VVC.

Accordingly, we asked whether susceptibility to VVC was evident in the absence of exogenous E2 in the Type 17-deficient mouse strains used in this study. In keeping with the well-described kinetics of murine VVC [15], WT controls given the sesame oil vehicle (SO, Veh) typically clear vaginal *C. albicans* by day 7 (**Fig 4a-d**). *Il23^-/-^* and *Il22ra1^-/-^* mice similarly cleared *C. albicans* by day 7 (**Fig 4a, b**), in line with prior findings that *Il17ra^-/-^* and *Il22^-/-^*individual knockout mice do not show elevated fungal loads without E2 [24]. PMN counts were the same in infected *Il23^-/-^* and *Il22ra1^-/-^*mice given vehicle compared to WT, though some variation was seen when comparing either knockout to E2-treated WT mice (**Fig S3B**).

**Figure 4.**
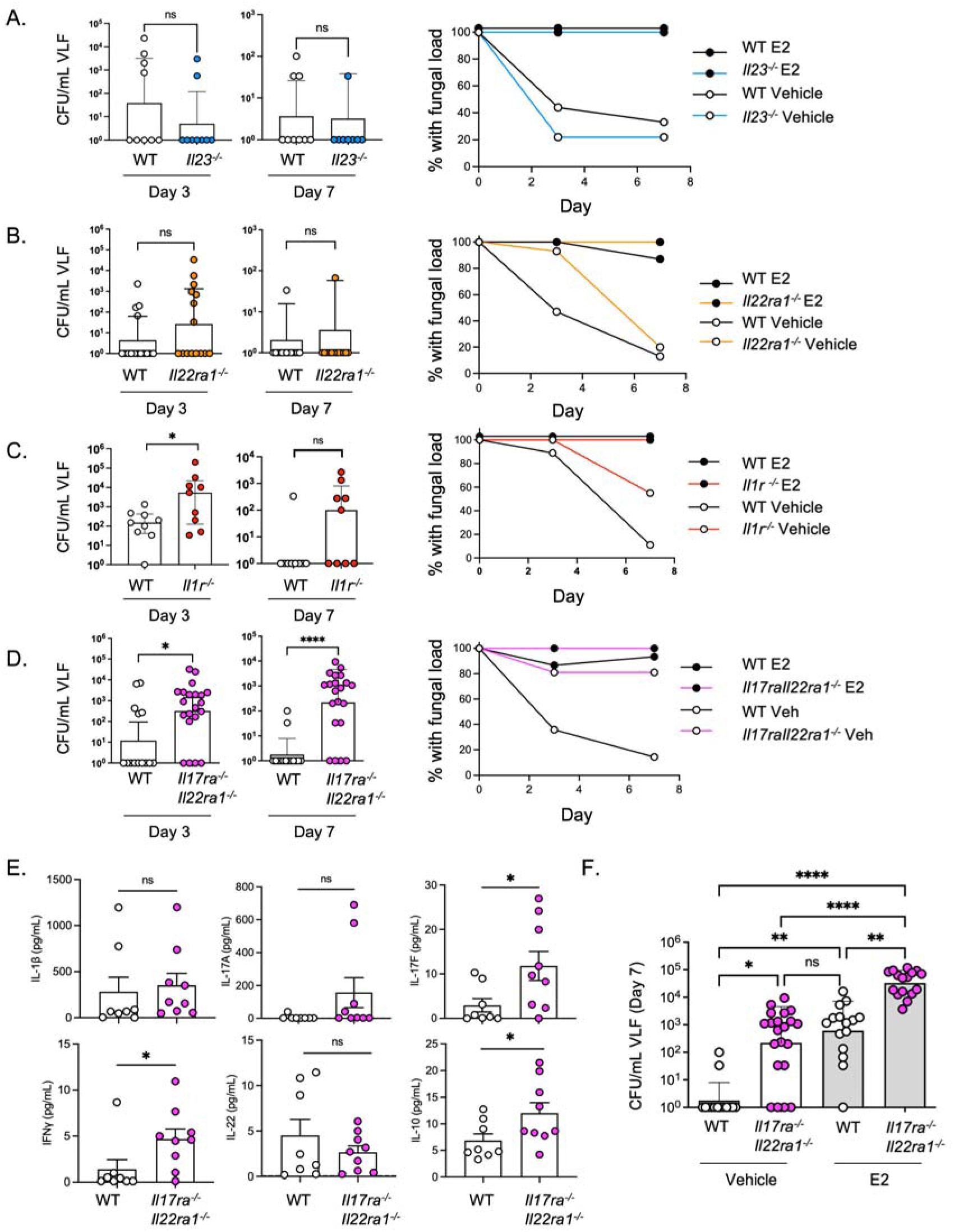
IL-17/22 receptor deficiency drives VVC susceptibility in the absence of E2 conditioning. **a-d.** Indicated mice were administered sesame oil (SO) vehicle (Veh). Left: fungal loads in VLF assessed on days 3 and 7 (Il23a^-/-^ n=8-9; Il22ra1^-/-^ n=10-11; Il1r^-/-^ n= 9; Il17raIl22ra1^-/-^ n=14-21). Right: Percent of mice in each cohort with a detectable fungal load at each time point. **e.** Cytokine levels in VLF on day 3. t-test with Welch’s correction. **f.** Direct comparison of fungal loads in E2-treated versus vehicle-treated Il17raIl22ra1^-/-^ mice (same experimental data as in panel **3a** and **4d**). Data in **a-d, f** show geometric mean ± geometric SD analyzed by two-tailed unpaired Mann-Whitney test.

However, non-estrogenized *Il1r^-/-^* and *Il17raIl22ra1^-/-^*mice both harbored higher fungal burdens and delayed clearance kinetics compared to WT controls (**Fig 4c, d**). PMN frequencies were not altered in *Il17raIl22ra1^-/-^* mice (**Fig S3C**). Unlike E2-treated VVC, *Il17raIl22ra1^-/-^*mice given SO vehicle did not display increased LDH activity in VLF (**Fig S3C**). Of the 40 cytokines and chemokines tested, only IFNγ and IL-17F were increased in VLF from these mice (**Fig 4e**, *data not shown*). Of note, vehicle-treated *Il17raIl22ra1^-/-^*mice had similar fungal burdens to estrogenized WT mice, implying that immune restriction and estrogen promotion of fungal colonization act in similar magnitudes (**Fig 4f**). E2 further elevated fungal loads in *Il17raIl22ra1^-/-^* mice, suggesting that these are additive pathways of susceptibility. Therefore, while IL-1/17/22 signaling governs control of fungal burden under both estrogen-free and - replete conditions, estrogen administration seemingly promotes an environment that facilitates elevated fungal-mediated damage.

E2 synchronizes the murine estrus cycle to a pseudoestrus phase, a state most receptive to fungal infection [63]. Although *Il17raIl22ra1^-/-^* mice are fertile with normal litter sizes, increased fungal loads in *Il17raIl22ra1^-/-^* mice could be explained by dysregulation of endogenous estrogen or the estrous cycle. To examine this, we tracked estrous cycle dynamics in WT and *Il17raIl22ra1^-/-^* mice over 3 cycles (12 days), collected at the same time of day to avoid circadian effects of hormones and IL-17 [64, 65] (**Fig 5a, b**). *Il17raIl22ra1^-/-^*and WT mice spent similar amounts of time in estradiol-high phases (proestrus/estrus) and estradiol-low phases (metestrus/diestrus) (**Fig 5c, d, Fig S5a**). Sera estradiol, cortisol, and testosterone levels were not different between WT and *Il17raIl22ra1^-/-^* mice at proestrus/estrus (**Fig 5e**). Unexpectedly, and despite their apparently normal estrous cycle dynamics, *Il17raIl22ra1^-/-^* mice had elevated progesterone (P4) levels in serum (**Fig 5e**). However, this is unlikely to explain their elevated fungal loads, since higher P4 concentrations would be expected to limit fungal colonization [13]. Since the opposite is seen in *Il17raIl22ra1^-/-^*mice, these data suggest that susceptibility to fungal colonization due to dual cytokine receptor deficiency is independent of steroid hormone status.

**Figure 5.**
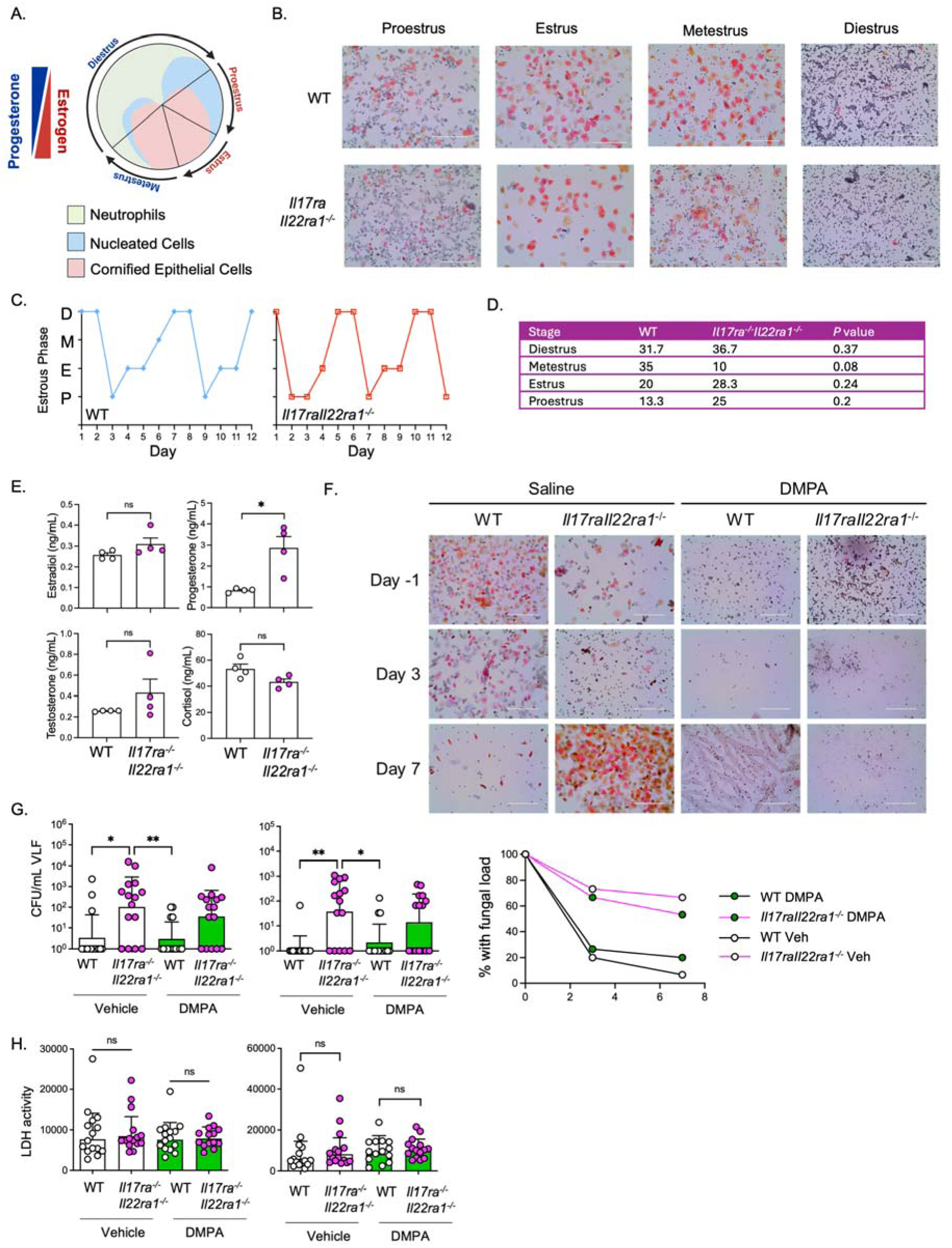
Sex hormone dynamics in Il17raIl22ra1^-/-^ mice. **a.** Hormone predominance and cell populations in VLF during the murine estrus cycle, adapted from Ref. [65] **b.** Sample PAP staining of VLF. **c, d.** Estrous phase distributions analyzed using per-mouse proportions and Wilcoxon rank-sum tests to account for repeated-measures **e.** Serum hormone levels at proestrus or estrus (n=4). two-tailed unpaired Student’s t-test **f.** Representative Pap smears after saline or DMPA treatment, assessed at day - 1, 3 and 7 relative to infection. **g.** Fungal loads in VLF on days 3 and 7. Saline (Vehicle) or DMPA was given on days -5 and 2 relative to infection All groups n=15. Mean ± geometric SD analyzed by two-tailed unpaired Mann-Whitney test. Right: Percent of mice in each cohort with a detectable fungal load at each time point. **h.** LDH activity in VLF on days 3 and 7. t-test with Welch’s correction

To ascertain if the Type 17-driven increase in fungal loads or tissue damage is hormone-driven, we administered mice depot medroxyprogesterone (DMPA), a progestin-based contraceptive, which reliably synchronized mice into durable pseudodiestrus throughout the duration of infection (**Fig 5f**). *Il17raIl22ra1^-/-^*mice given DMPA had higher fungal loads compared to WT (**Fig 5g**), although tissue damage as measured by LDH was not altered (**Fig 5h**). DMPA is not linked to increased VVC in humans [66], and prior reports showed that administration of progesterone (P4) in mice does not enhance *C. albicans* vaginal colonization [13], but may actively promote fungal clearance [63, 67, 68]. Consistent with this, we found that *Il17raIl22ra1^-/-^*mice given P4 trended to higher fungal loads compared to WT (**Fig S5D**), with the caveat that P4 was not as effective as DMPA in locking mice out of estrus. P4-treated *Il17raIl22ra1^-/-^* mice also exhibited increased LDH activity compared to P4-treated WT mice, unlike DMPA-treated mice (**Fig S5E**), likely because a subset of mice were still cycling.

Collectively, these data support the existence of a substantial Type 17-dependent contribution to antifungal control in the vaginal tract that is operative regardless of estrous cycle stage. Moreover, Type 17 loss does not detectably impact the estrous cycle, levels of estrogen, or fertility.

## Discussion

An estimated 75% of childbearing aged women develop VVC at least once in their lifetime. As many as 5-8% suffer from recurrent infections, emphasizing a need to better understand the underlying pathogenesis of this condition for development of targeted therapeutic interventions [4, 7, 69]. *C. albicans* infects distinct anatomical tissues with quite different immune susceptibility profiles, and thus inferences from studies in the mouth, gut or visceral organs do not necessarily apply in the vaginal mucosa. Numerous studies have described genetic polymorphisms associated with VVC, many of which identified pattern recognition receptors and innate-acting immune components [70, 71]. However, as noted, there is considerable evidence for an adaptive immune component in VVC. Viewed within the ‘damage-response framework’, VVC is a function of immune pathology arising from host immunity as well as a failure to control the fungus [8, 10].

Our data indicate that IL-1/Type 17 immunity in VVC mediates colonization resistance and mitigates estradiol-related tissue damage that occurs in VVC. There is a stronger correlation of tissue damage in the absence of Type 17 immunity which suggests that *C. albicans* is more damaging in a setting lacking IL-17R/IL-22R, consistent with findings in OPC, where loss of IL-17 or IL-22 is associated with increased hyphal formation and candidalysin-induced epithelial damage [39, 48, 72–75]. The Type 17 pathway is operative even in the absence of the estrogenic pathways that are a predominant driver of disease. Thus, VVC can be viewed as arising from independently-acting routes. One pathway to VVC susceptibility is driven by estradiol sensitization of the vaginal microenvironment and/or effects of estrogen on *C. albicans* itself [4, 10]. An alternative path is through an IL-1R-dependent IL-17/IL-22-driven cytokine response. Importantly, loss of both susceptibility pathways leads to more fulminant disease than loss of either pathway alone. This conclusion is supported by the observation that fungal loads are the same in E2-treated WT mice compared to *Il17raIl22ra1^-/-^* mice, indicating both routes contribute equivalently to establishing disease. E2 administration combined with IL-17R/22R-deficiency or IL-1R-deficiency rendered mice more susceptible than either condition alone, indicating in either setting (E2 or cytokine deficiency) the other pathway is still operative.

IL-1 cytokines are highly pleiotropic, and IL-1R signals drive innate and adaptive responses to candidiasis, including Type 17-driven immunity in OPC [75–78]. However, its ability to promote Type 17 immunity in VVC was not thought to be a likely mechanism of action since neither IL-17 nor IL-22 individually played a prominent role in VVC [24, 26, 31, 35]. IL-1 acts upstream of Type 17 responses in vaginal immunity to herpes simplex virus and chlamydial infections [42, 43, 75, 76, 79, 80], so our findings are in line with other studies of IL-1R function in the reproductive tract. There is not compelling evidence linking clinical IL-1 inhibition in humans to VVC, which could be due to underreporting or differences in how the Th17 axis is controlled in mice versus humans [81–83].

The essential role of IL-17 in oral and dermal candidiasis has been recognized for over 15 years [84], but its contribution to VVC immunity has been less obvious. Studies of mice lacking individual genes in these pathways surprisingly, did not reveal contributions of IL-17A/F, Act1, or IL-22 signaling [24, 31]. Similarly, rare individuals with inborn mutations in Th17-related cytokines are highly prone to mucocutaneous oral and dermal candidiasis but not VVC [85–87]. VVC is also not apparent in patients taking anti-IL-23 biologics [26], consistent with findings in *IL23*^-/-^ mice (this study, [31]). Anti-IL-22 Abs are not approved for clinical use, so its contributions to clinical VVC in humans remains unknown. *AIRE* mutations causing the autoimmune syndrome APECED are accompanied by anti-IL-17A/F and anti-IL-22 Abs, sometimes contemporaneously, but VVC is not a hallmark of this syndrome [88]. Historically then, these data suggested that immunity to VVC, unlike most manifestations of candidiasis, was not driven by Th17 cells.

Despite this, we were prompted to revisit the possibility that combinatorial activities of IL-17 and IL-22 in VVC susceptibility. There is a potent induction of IL-17 and IL-22 in the reproductive tract during VVC [23, 24]. A meta-analysis of over 17,000 patients taking anti-IL-17 biologics for autoimmune diseases reported a modest but statistically significant increase in VVC incidence [26]. OPC is far more severe in *Il17raIl22ra1^-/-^* mice compared to mice lacking either receptor alone [48], hinting that the antifungal properties of these cytokines are potentially synergistic. CARD9-deficient mice showed impaired fungal control in VVC, associated with reduced Type 17 cytokine responses [89]. We found that a human vulvar cell line widely used in the field to study vaginal responses to *C. albicans* showed potent synergistic responses to IL-17 and IL-22 in combination, though responses to either cytokine alone were modest. Thus, although loss of either IL-17 or IL-22 signaling alone can be tolerated in the vaginal mucosa in terms of antifungal immunity, loss of both causes high and persistent colonization.

Our data reveal a divergence of IL-1R versus Type 17 signals with respect to control of neutrophils. Whereas *Il1r*^-/-^ mice showed modestly impaired PMN numbers, *Il17raIl22ra1^-/-^*mice trended to increased PMNs, concomitant with high fungal loads. Increased vaginal fungal burdens and reduced neutrophil influx in *Il1r^-/-^* mice were similarly reported in murine OPC, where keratinocyte-derived IL-1[Z] contributed to neutrophil recruitment [77]. Thus, IL-1 signaling appears to control distinct arms of VVC immunity – one pathway activates PMNs which are insufficient to contain disease, and the other activates Type 17 cytokines which mediate immune control. While IL-1R signaling is driven by both IL-1[Z] and IL-1β, our data indicate that IL-1β dominates for control of vaginal fungal burden, though surprisingly not the PMN response. Thus, delineating isoform- and compartment-specific roles of IL-1 isoforms is warranted.

In light of the susceptibility of *Il1r*^-/-^ mice but not *Il23*^-/-^ mice to VVC, it appears that IL-1R signaling plays a dominant role in shaping the Type 17 cytokine response within the vaginal tract. The source of cytokines was not examined in this study. IL-1 cytokines, like their receptors, are broadly expressed. Vaginal epithelial cells produce high levels of IL-1 cytokines in response to *C. albicans* [5, 6, 90, 91]. Unlike humans, mice are immunologically naïve to *C. albicans* [50, 75, 92, 93], so given that effects in the VVC model are observed as early as day 3, before antigen-specific adaptive immunity is established, it is likely that producers of IL-17A/F and IL-22 are innate-acting lymphocytes, such as γδ-T cells [94–102]. Much like the Type 17 pathway, the contributions of γδ-T cells to host defense during VVC is not fully resolved [95, 96, 103].

Multiple fungal factors contribute to VVC and influence host cytokine responses. Candidalysin (encoded by *Ece1*) is a secreted pore-forming fungal toxin required to induce the Type 17 response in OPC via epithelial tissue damage [75, 76, 104] and promotes tissue damage in VVC [5, 38]. Pra1, which facilitates zinc metabolism in *C. albicans*, is associated with the elevated cytokines that characterize VVC pathology [105]. There is a strong influence of vaginal microbiota in VVC [69, 106], which could be altered in *Il17raIl22ra1^-/-^* mice.

VVC differs from other manifestations of candidiasis in being hormone-dependent. High-estrogenic oral contraceptives, hormone replacement therapies, and pregnancy all increase the risk of VVC [7, 69]. Still, the mechanism for E2-driven susceptibility is not well understood. Estrogen is thought to act on the host through the Esr1 receptor [68], potentially by creating an immunosuppressive environment [107]. Estrogen also impacts *C. albicans* directly, promoting its virulence and reducing immune recognition [59, 61]. E2 is also implicated in systemic candidiasis through impairing IL-23 production in DCs [108], presenting an intriguing contrast to VVC where IL-23 is not involved. Since most prior studies of gene-targeted mice have not included non-estrogenized controls, the potent susceptibility caused by estrogen condition may have masked contributions from other arms of immunity in this setting.

A surprise from this work was that *Il1r*^-/-^ and *Il17raIl22ra1^-/-^* mice exhibited substantially elevated vaginal fungal loads even in the absence of exogenous E2. Increased VVC susceptibility occurred in *Il17raIl22ra1^-/-^* mice despite their abnormally elevated progesterone seen at baseline. However, progesterone would not be predicted to increase *C. albicans* susceptibility; in fact, *Il17raIl22ra1^-/-^*mice retained high fungal loads even under the influence of DMPA or P4. Progesterone-driven restriction of VVC has been linked to TNF and neutrophils [63, 67, 68], and our data suggest that IL-17R/IL-22R signaling contributes in this setting as well. It is not straightforward to test the impact of estrogen directly, as estrogen receptor inhibitors such as tamoxifen have direct antifungal activity [109], and ovariectomy procedures are invasive with broad physiological effects.

As with all animal models, VVC in mice has limitations [110]. The murine model does not mirror RVVC, where most human genetic studies have been performed [7, 71]. The mouse estrous cycle and vaginal pH are quite different [65], as are microbial communities [111, 112]. The role of IL-1 in driving Th17 differentiation is not identical in mice and humans [53]. Still, with respect to IL-17/Th17 cells and *C. albicans* infections, scrutiny has revealed consistent similarities with respect to impact of gene impairment, antigenic responses, and contributions of adaptive immunity.

Whereas loss of either IL-17 or IL-22 signals alone are insufficient to cause hallmark VVC symptoms, we find that loss of both elicits not only a profound elevation in fungal colonization but also tissue damage and inflammation. Estrogen-driven susceptibility to VVC is a key driver of tissue damage, distinguishing it from other manifestations of candidiasis. These data show that immunity to VVC is also controlled by an IL-1, IL-17, and IL-22 cytokine circuit that acts independently of the hormonal regulation of fungal colonization. Collectively, these data may reconcile some of the disparate findings in the field with regards to VVC and the Type 17 pathway.

## Materials and Methods

### Candida albicans and VVC

One colony of *Candida albicans* SC5314 (ATCC) was grown overnight in Yeast Peptone Dextrose (YPD) broth with 50 μg/mL chloramphenicol at 30°C. 5×10^6^ CFU *C. albicans* yeast in PBS was administered by intravaginal inoculation [24]. If indicated, on days -3 and 4 relative to infection, mice were subcutaneously administered 0.1 mg of β-estradiol 17-valerate (E2; Millipore-Sigma, E1631) or 2 mg progesterone (MilliporeSigma, P0130) in SO vehicle. For DMPA studies, mice were given 0.9% NaCl (saline) vehicle or 2 mg DMPA in saline on days -5 and 2 relative to infection. At days 3 and 7, VLF was collected by PBS wash (100 µL) and colonies enumerated on YPD-agar.

### Mice

Animal protocols were approved by the University of Pittsburgh and the University of Tennessee Health Science Center Institutional Animal Care and Use Committees. Wild-type (WT) mice were from the Jackson Laboratory, Taconic Farms, or generated in-house from breeding. *Il1r*^-/-^(strain #003245) mice were from The Jackson Laboratory and *Il1b*^-/-^ mice were from N. Archer [113]. *Il23a*^-/-^ mice were from Genentech and *Il17ra*^-/-^ mice were from Amgen. *Il22ra1Il17ra^-/-^*mice were described and were fertile with normal litter sizes [48]. Mice were housed in SPF conditions and used at 6-12 weeks of age. All mice were on the C57BL/6 background.

### Cell Culture

A-431 epithelial cells (ATCC, CRL-1555) were cultured in DMEM with 10% FBS and 1% penicillin/streptomycin and stimulated with human IL-17A (100 ng/mL) or IL-22 (100 ng/mL) (Peprotech) in replete media.

### Microscopy

PMN recruitment was assessed in VLF smears by Papanicolaou (PAP) staining. PMNs were enumerated in 5 nonadjacent fields on an EVOS FL Auto Imaging system (Invitrogen) acquired at 20X or 40X and counts normalized to a 20X field. Investigators were blinded to sample identity. Data presented as the average PMN count from 5 images. Excised vaginal tissue was fixed in 10% neutral buffered formalin, paraffin-embedded, and slides were co-stained with Periodic acid–Schiff (PAS), hematoxylin, and eosin and imaged on an EVOS FL Auto Imaging system (Invitrogen). Size bars included in images.

### Estrous phase tracking, hormone and blood glucose measurement

VLF was collected at approximately the same time daily for 12 consecutive days. Estrous cycle phase was assessed by Pap staining [65]. For hormone assessment, blood from mice proestrus/estrus or estrus was collected by cardiac puncture. Sera was isolated using serum separator tubes (BD Microtainer, 365967). Hormones were measured using the Steroid/Thyroid 6-Plex Assay Multi Species Array panel (Eve Technologies Corporation, Calgary, Canada). Blood glucose was measured from tail nicks in naïve, unfasted mice using a Contour Next EZ glucometer and Countour Next test strips.

### RNASeq, qPCR

RNA from frozen vaginal tissue or A-431 cells was isolated using RNeasy Mini Kits (Qiagen) and cDNA generated with the iScript cDNA Synthesis Kit (Bio-Rad). Real-time qPCR was performed using 2X Universal SYBR Green Fast qPCR Mix (ABclonal) on a CFX96 Real-Time PCR Detection System (Bio-Rad). Quantitect primer assays were used for transcript quantification (Qiagen). Sequencing data is registered with the Bioproject database (PRJNA1245436) and will be made available upon paper acceptance.

### ELISA, Luminex, LDH

Cytokines and chemokines were measured in clarified VLF harvested on day 3 using Luminex xMAP technology by Eve Technologies Corporation (Calgary, Canada) with the mouse Cytokine/Chemokine 32-Plex Discovery Assay Array (MD32), the Mouse Cytokine/Chemokine 36-Plex Discovery Assay (MD36) or the Cytokine Th17 12-Plex Discovery Assay Array (MDTH17-12). For samples too dilute for calculation, values were plotted at the manufacturer-specified limit of detection for each analyte. LDH in VLF was measured using CytoTox 96 Non-Radioactive Cytotoxicity Assay (Promega, G1780). OD_492_ was determined using a BioTek Synergy 2 microplate reader. Samples were quantified against a standard curve of purified LDH (Sigma, L7525).

### Statistics

Data were analyzed using Prism (GraphPad Software Inc.) as described in figure legends. Data are pooled from at least two independent experiments unless noted. Throughout, *P < 0.05, **P < 0.01, and ***P < 0.001; ns, not significant.

## Supporting information

Merged supplementary data

## Acknowledgments

SLG was supported by the University of Pittsburgh and NIH (AI185365). TCT and MEC were supported by T32-AI089443, and MEC by F32-AI186291. BP was supported by AI134796. The University of Pittsburgh Biospecimen Core was supported, in part, by the University of Pittsburgh Office of the Senior Vice Chancellor for Health Sciences. The funders had no role in study design, data collection and analysis, decision to publish, or preparation of the manuscript. We thank Amgen for *Il17ra^-/-^* mice and Genentech for *Il23a*^-/-^ mice. *Il1b*^-/-^ mice were a gift from N. Archer (Johns Hopkins) and supplied by Y Iwakura (University of Tokyo). We thank C. Kausic and M. Bertolini for valuable input.

## Abbreviations

CFU, colony forming units; E2, 17-β-estradiol; DMPA, depot medroxyprogesterone; OPC, oropharyngeal candidiasis; LDH, lactate dehydrogenase; P4, progesterone; Pap, Papanicolaou; SO, sesame oil; VVC, vulvovaginal candidiasis; Veh, vehicle; VLF, vaginal lavage fluid

---

**Fig S1.**
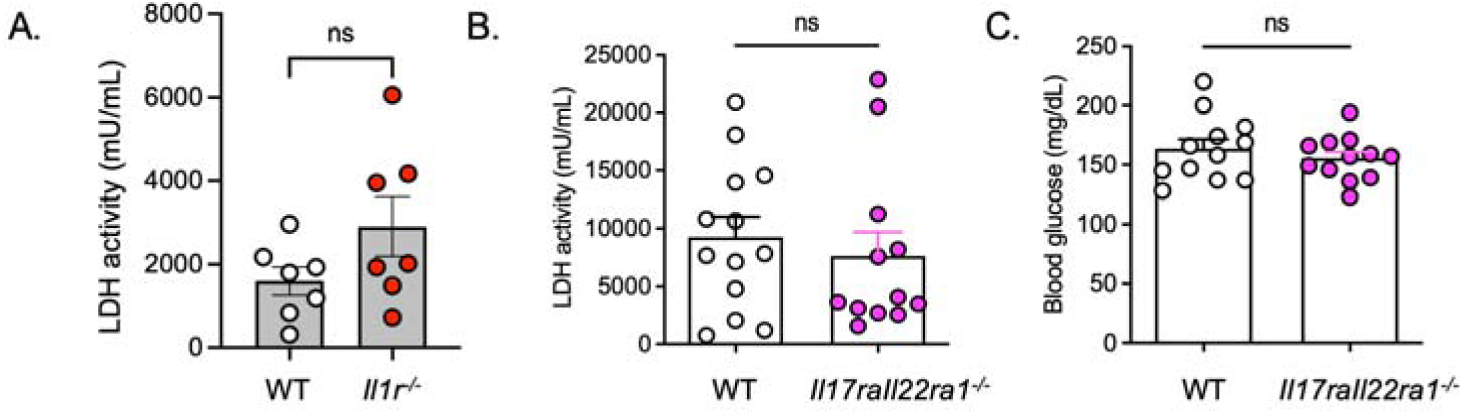
Baseline LDH activity and blood glucose levels in Type 17-deficient mice. **a, b.** LDH levels in VLF (mU/mL) on day 3. Mean+SEM, analyzed by t-test with Welch’s correction. **c.** Blood glucose levels in serum (mg/dL). Mean+SEM, Welch’s correction

**Fig S2.**
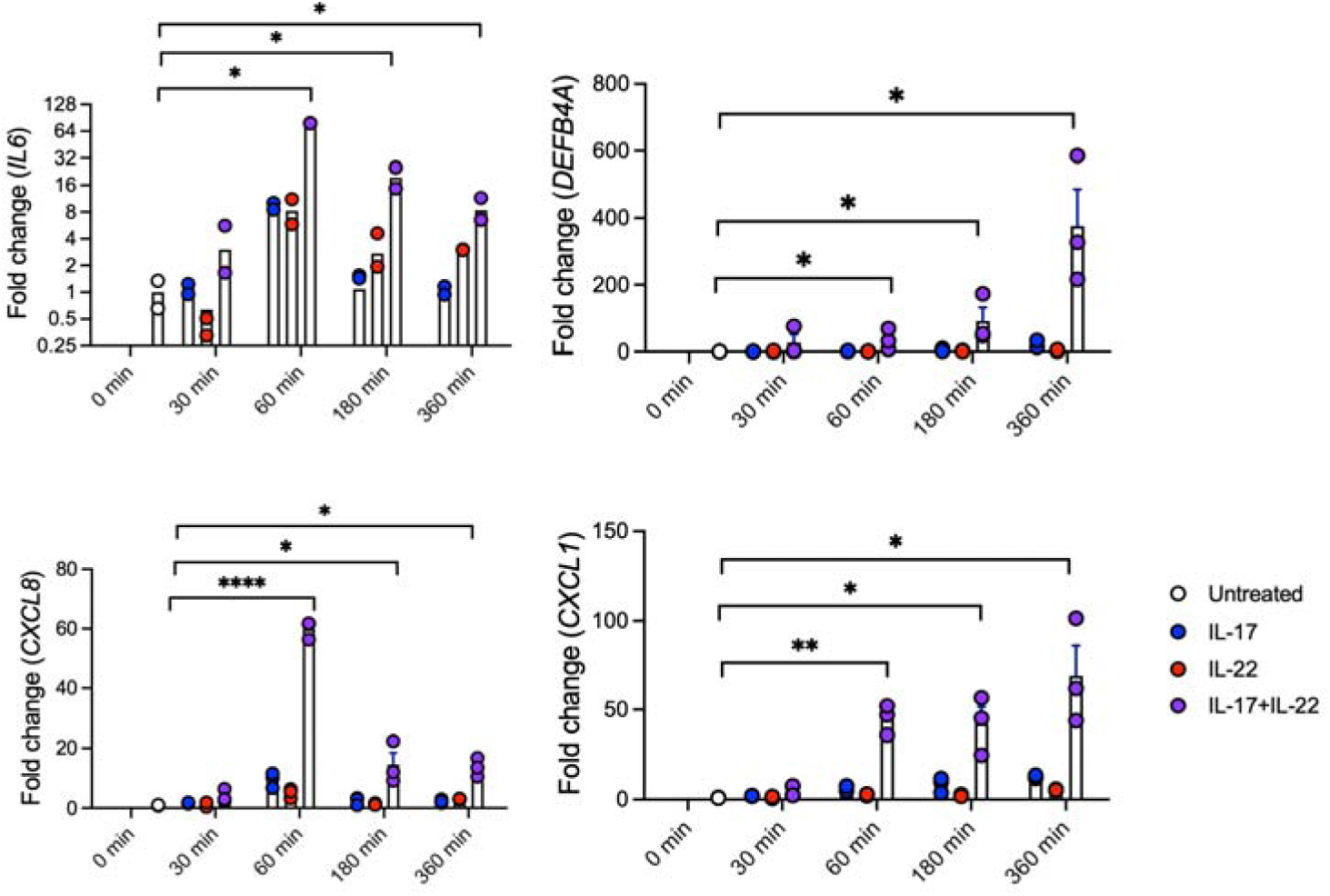
**IL-17 and IL-22 synergy in human vaginal epithelial cells**. A-431 vulvar epithelial cells were treated with human IL-17 and/or IL-22 for the indicated times. Genes were assessed by qPCR relative to GAPDH and normalized to untreated control (0 minutes). Mean+SEM, analyzed by ANOVA and Tukey’s multiple comparisons test

**Fig S3.**
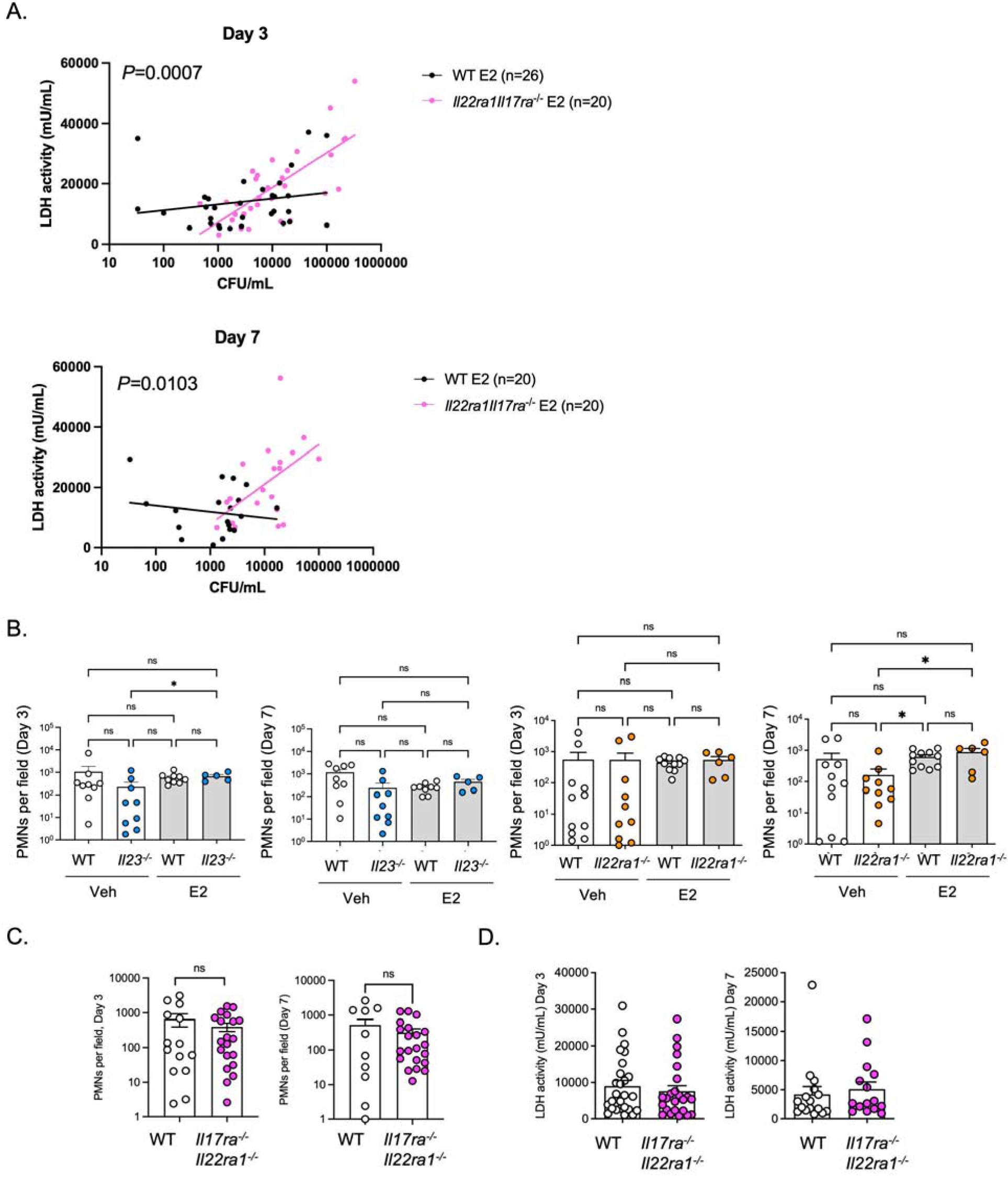
VVC parameters in gene targeted mice. **a.** Correlation of LDH and CFU levels in estradiol (E2)-treated WT or Il17raIl22ra1^-/-^ mice. Slopes compared by nonlinear regression. **b.** PMN counts in IL-23^-/-^ and IL-22ra1^-/-^ mice. PMNs in VLF determined by cytology, analyzed by two-tailed unpaired Student’s t-test with Welch’s correction. **c.** PMN counts on day 3 in SO treated Il17raIl22ra1^-/-^ mice on dyas 3 and 7. Mean+SEM, analyzed by student’s t-test with Welch’s correction. **d.** LDH activity in VLF on day 3 and 7. Mean+SEM, analyzed by student’s t-test.

**Fig S4.**
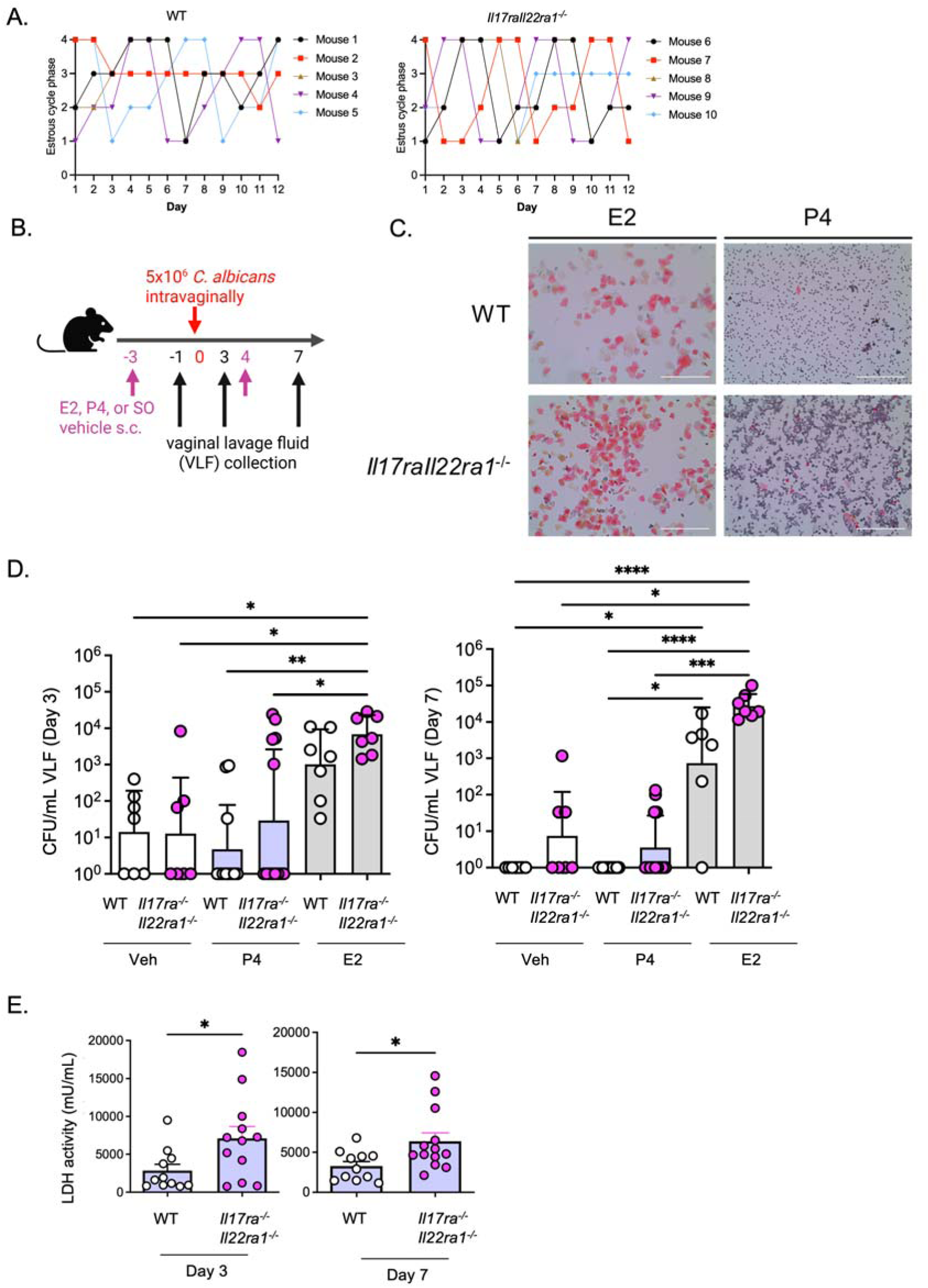
**a**. Estrous cycle tracking by PAP staining of VLF over 12 days. D: diestrus, M: metestrus, E: estrus, P: proestrus/estrus. **b**. Experimental setup**. c**. Representative PAP staining of day -1 VLF after E2 or P4 treatment. **d.** Fungal loads in VLF on days 3 and 7 after sesame oil (Vehicle) (n=7), estrogen (E2) (n=6-7), or progesterone (P4) (n=11-13) given on days -3 and 4 relative to infection. Mean ± geometric SD analyzed by two-tailed unpaired Mann-Whitney test. **e.** LDH activity in VLF on days 3 and 7 in P4-treated mice. t-test with Welch’s correction

