## Supplementary material for "An IL-1, IL-17, and IL-22 cytokine circuit controls vulvovaginal candidiasis independently of estrogen": Merged supplementary data

*Short title: VVC is controlled by immune cytokines independently of hormones*

Bianca M. Coleman[1*], Melissa E Cook[1*], Md. Robin Khan[2], Amanda K. Vogel[3], Anthony J. Wells[4], Jian Miao[2], Shachi P. Vyas[1], Tiffany C. Taylor[1], Felix E.Y. Aggor[1], Nicole O. Ponde[1], Ipsita Dey[1], Henry Zou[1], Eldin Jašarević[5-7], Brian M. Peters[4¶*], Sarah L. Gaffen[1¶*]

[1] Division of Rheumatology & Clinical Immunology, University of Pittsburgh, Pittsburgh PA 15261, USA

[2] Graduate Program in Pharmaceutical Sciences, College of Graduate Health Sciences, University of Tennessee Health Science Center, Memphis TN 38163

[3] Integrated Program in Biomedical Sciences, College of Graduate Health Sciences, University of Tennessee Health Science Center, Memphis TN 38163

[4] Department of Clinical Pharmacy and Translational Science, College of Pharmacy, University of Tennessee Health Science Center, Memphis TN 38163

[5] Department of Obstetrics, Gynecology and Reproductive Sciences, University of Pittsburgh, Pittsburgh PA 15213

[6] Department of Computational and Systems Biology, University of Pittsburgh, Pittsburgh PA 15213

[7] Magee-Womens Research Institute, University of Pittsburgh, Pittsburgh PA 15213

** These authors contributed equally to this manuscript*

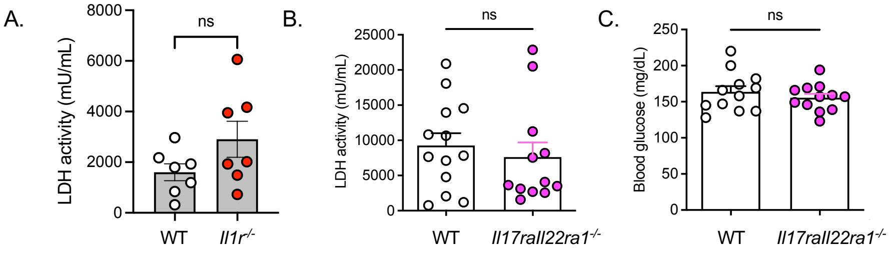


**Fig S1. Baseline LDH activity and blood glucose levels in Type 17-deficient mice. a, b.** LDH levels in VLF (mU/mL). Mean+SEM, analyzed by t-test with Welch's correction. **c.** Blood glucose levels in serum (mg/dL).

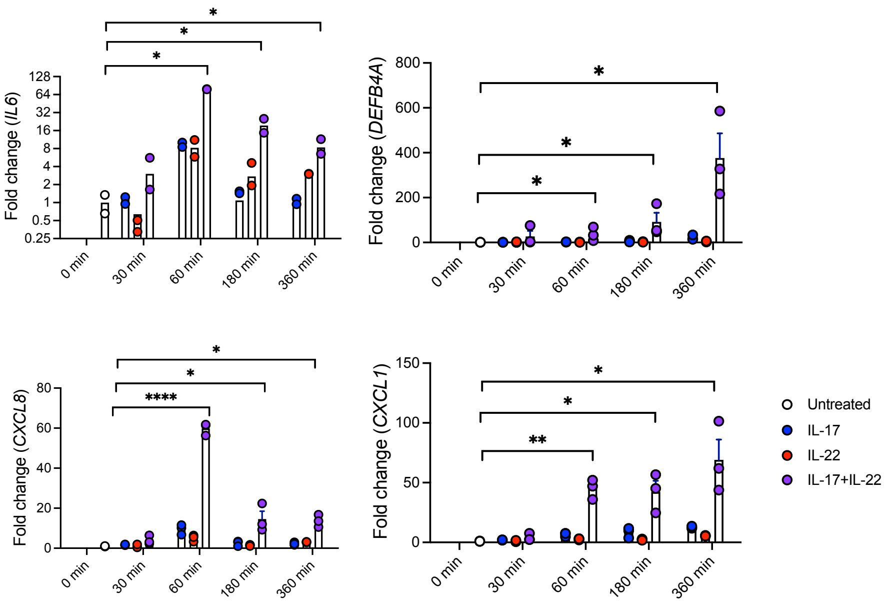


**Fig S2**. **IL-17 and IL-22 synergy in human vaginal epithelial cells**. A-431 vulvar epithelial cells were treated with human IL-17 and/or IL-22 for the indicated times. Genes were assessed by qPCR relative to GAPDH and normalized to untreated control (0 minutes). Mean+SEM, analyzed by ANOVA and Tukey's multiple comparisons test

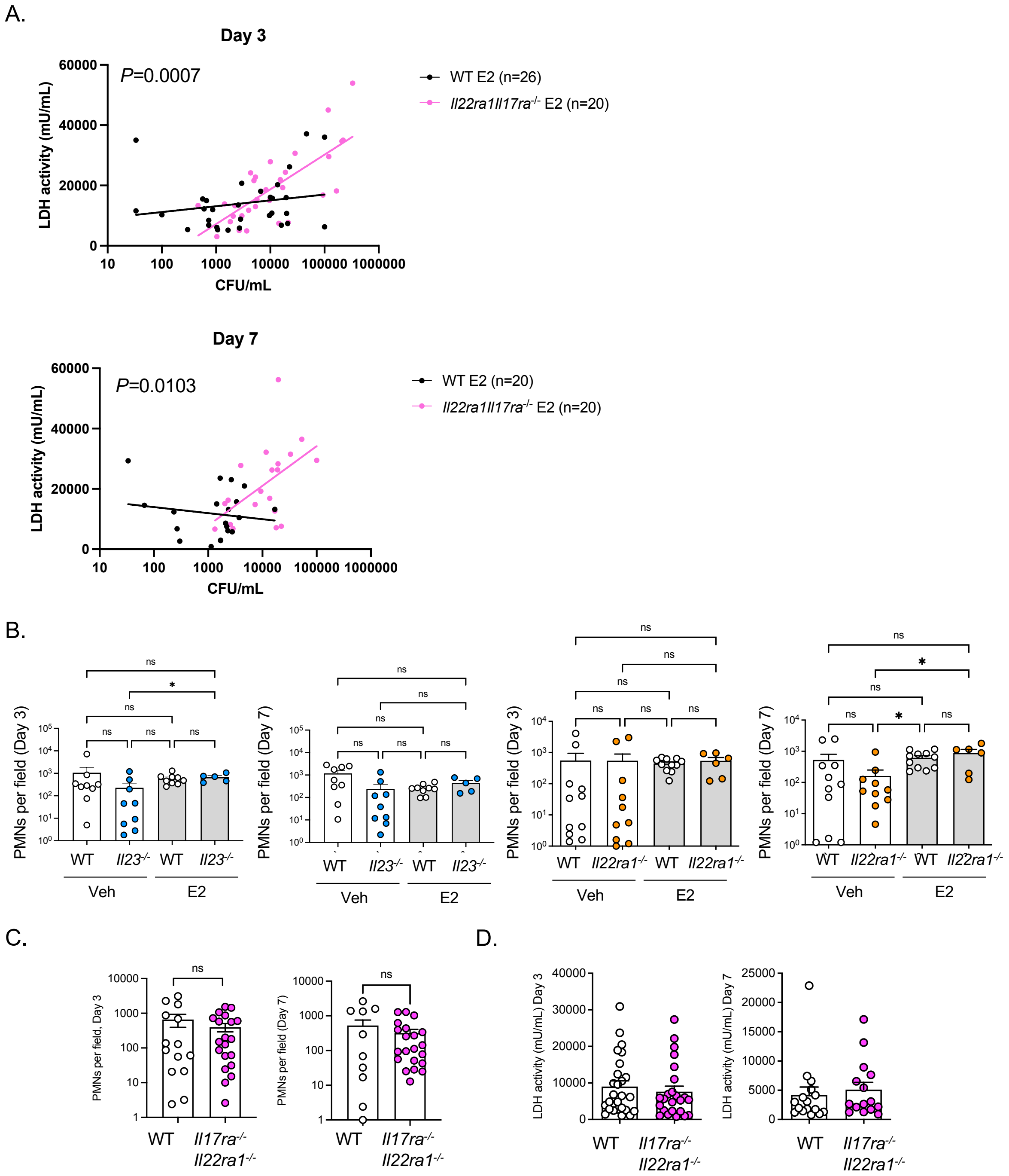


**Fig S3. VVC parameters in gene targeted mice. a.** Correlation of LDH and CFU levels in estradiol (E2)-treated WT or *Il17raIl22ra1*$^{-/-}$ mice. Slopes compared by nonlinear regression. **b.** PMN counts in *Il23*$^{-/-}$ and *Il22ra1*$^{-/-}$ mice treated with SO vehicle or E2. PMNs in VLF determined by cytology. Mean+SEM, analyzed by two-tailed unpaired Student's t-test with Welch's correction. **c.** PMN counts on day 3. Mean+SEM, analyzed by student's t-test with Welch's correction. **d.** LDH activity in VLF on day 3 and 7. Mean+SEM, analyzed by student's t-test.

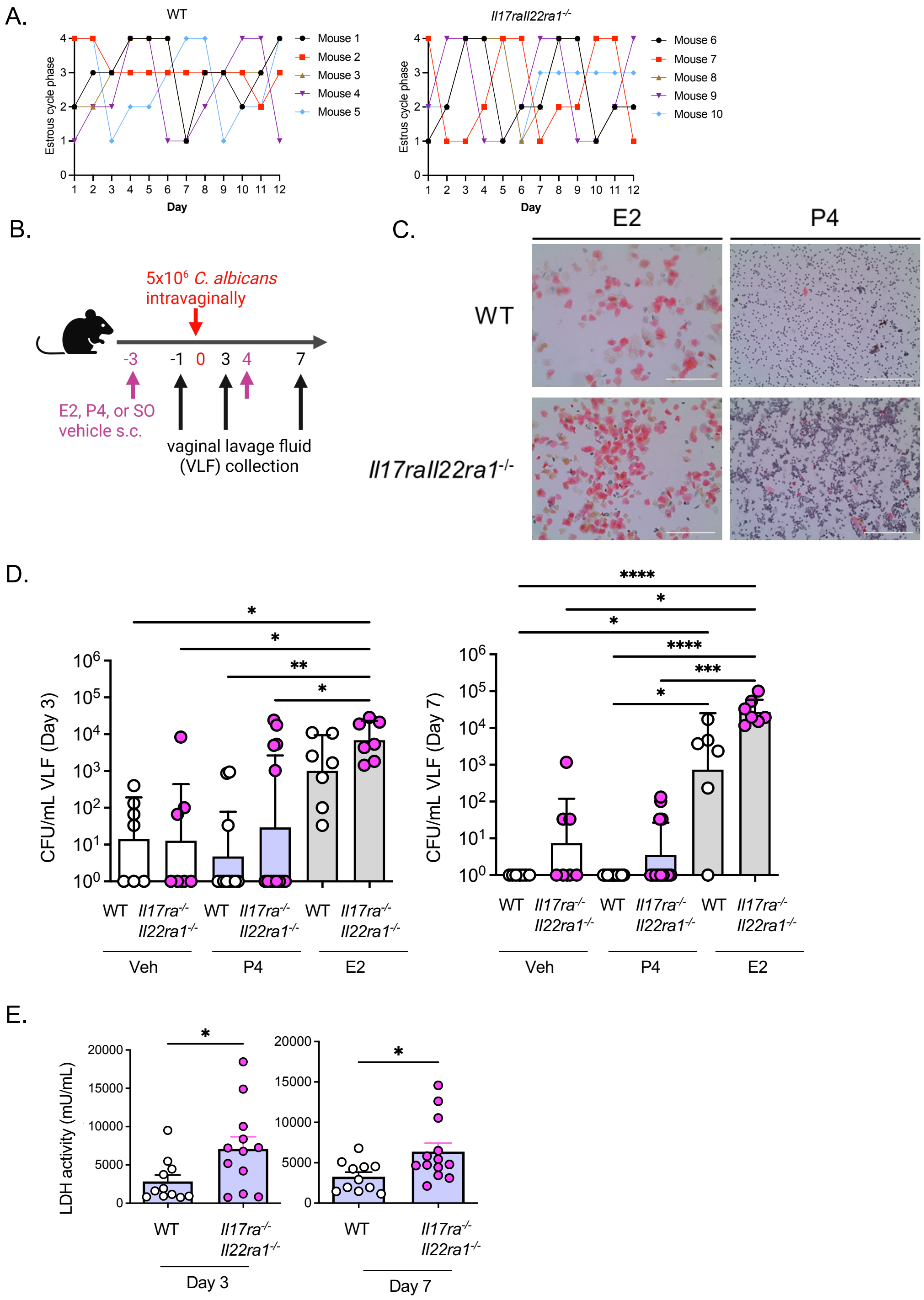


**Fig S4. Progesterone does not alter susceptibility to VVC**. **a**. Estrous cycle tracking by PAP staining of VLF over 12 days. D: diestrus, M: metestrus, E: estrus, P: proestrus/estrus. **b**. Experimental setup**. c**. Representative PAP staining of day -1 VLF after E2 or P4 treatment. **d.** Fungal loads in VLF on days 3 and 7 after sesame oil (Vehicle, n=7), estrogen (E2, n=6-7), or progesterone (P4, n=11-13) given on days -3 and 4 relative to infection. Mean ± geometric SD analyzed by two-tailed unpaired Mann-Whitney test. **e.** LDH activity in VLF on days 3 and 7 in P4-treated mice. t-test with Welch's correction
